# DesiRNA: structure-based design of RNA sequences with a Monte Carlo approach

**DOI:** 10.1101/2023.06.04.543636

**Authors:** Tomasz Wirecki, Grzegorz Lach, Farhang Jaryani, Nagendar Goud Badepally, S. Naeim Moafinejad, Gaja Klaudel, Janusz M. Bujnicki

**Affiliations:** Laboratory of Bioinformatics and Protein Engineering, International Institute of Molecular and Cell Biology in Warsaw, ul. Ks. Trojdena 4, 02-109 Warsaw, Poland; Institute of Theoretical Physics, Faculty of Physics (FUW), University of Warsaw, ul. Hoża 69, 00-681 Warsaw, Poland

**Author notes:** To whom correspondence should be addressed.; Correspondence may be also addressed to Tomasz Wirecki.

## Abstract

RNA sequences underpin the formation of complex and diverse structures, subsequently governing their respective functional properties. Despite the pivotal role RNA sequences play in cellular mechanisms, creating optimized sequences that can predictably fold into desired structures remains a significant challenge. We have developed DesiRNA, a versatile Python-based software tool for RNA sequence design. This program considers a comprehensive array of constraints, ranging from secondary structures (including pseudoknots) and GC content, to the distribution of dinucleotides emulating natural RNAs. Additionally, it factors in the presence or absence of specific sequence motifs and prevents or promotes oligomerization, thereby ensuring a robust and flexible design process. DesiRNA utilizes the Monte Carlo algorithm for the selection and acceptance of mutation sites. In tests on the EteRNA benchmark, DesiRNA displayed high accuracy and computational efficiency, outperforming most existing RNA design programs.

## INTRODUCTION

Ribonucleic acid (RNA) molecules represent integral components of biological systems, fulfilling crucial functions across a multitude of biological processes in living organisms. From transmitting genetic information from DNA to proteins, to sensing and communicating responses to cellular and extracellular signals, to catalyzing chemical reactions, the versatile functionality of RNA is indisputable (1). RNA molecules can also be used as potent biotechnology instruments, finding wide-ranging applications that span the development of RNA-based vaccines and therapeutics, the construction of biosensors, gene regulators, and components of the gene-editing machinery. The molecular and cellular activities of an RNA molecule often necessitate it to fold into a specific structure to perform its function. The RNA sequence determines the formation of canonical base pairs that form the secondary structure, and subsequently contribute to the formation of a three-dimensional structure of the molecule. Some RNA molecules form unique structures, while others form alternative structures or undergo transitions between the structured and unstructured states (2).

Various computational tools have been developed that predict RNA secondary structure from sequence (3). Two main families of programs can be distinguished: single-sequence and comparative methods. A classical approach for predicting RNA structure from a single sequence relies on the assumption that the native RNA structure is the one with the minimum free energy (MFE). The prediction is often attained by calculating the combination of all possible base pairs, implemented in methods using dynamic programming, which are deterministic by nature and guarantee finding the structure with the lowest free energy [e.g. as in RNAfold (4)]. A challenge connected to exploring all possible conformations of a given RNA molecule is that any RNA of length n can form 1.8^*n*^ possible secondary structures (5).

The complexity of the RNA folding problem is often reduced by introducing heuristics to identify a structure that is likely to be biologically relevant. One class of modern methods sample structures from the Boltzmann ensemble to identify large clusters of similar secondary structures with low (suboptimal) energies (6,7). Another class of methods use statistical models [e.g., as in Contrafold (8)] or machine learning [e.g., as in ContextFold (9)] without directly relying on physics-based models of RNA folding. Yet another class of methods infer the prediction based on evolutionary considerations. They require a multiple alignment of RNA sequences related to the target sequence. The most successful methods combine the comparative prediction with the minimum free energy structure calculations [e.g. as in PETFold (10)]. It is worth emphasizing that base-pairing may define not only secondary structure in the simplest sense, but also tertiary interactions such as pseudoknots and kissing loops. While many of the above-mentioned programs ignore such interactions for the sake of computational speed, there are algorithms that specialize in predictions for potentially pseudoknotted structures, such as IPknot (11). The relative predictive power of various methods for RNA secondary structure prediction on various datasets, for RNAs of various length, and with or without pseudoknots has been tested in the CompaRNA experiment (3) and in many other benchmarks.

Understanding the molecular basis of RNA functions, especially those extending beyond protein coding, requires an in-depth understanding of RNA structure [review (12)]. Consequently, one of the grand challenges that synthetic biology faces is the computational design of RNA molecules with desired spatial structures and functional attributes. The significance of this challenge is reflected in the numerous applications that RNA design could enhance, ranging from the development of improved RNA vaccines, modifying HIV-1 replication mechanisms (13) and reprogramming cellular behavior (14), to developing novel logic circuits (15) and constructing intricate RNA nano-objects (16). Moreover, artificially designed RNA sequences can be exploited as tools for the detection of novel naturally occurring RNAs (17) and other molecules.

The computational design of RNA calls for solutions to the so-called RNA inverse folding problem, defined as designing a sequence that folds into a predetermined target secondary structure under specific conditions. The complexity of this problem is exacerbated by the exponential growth in the number of possible sequences matching a given structure (6^*p/2*^4^*u*^ for an RNA having respectively u unpaired and p paired residues) (18,19), which renders a brute force search strategy impractical. Instead, efficient search strategies usually involve mutating a seed sequence iteratively and predicting the secondary structure of each variant until a sequence is obtained that matches the target structure.

The seminal work of Dirks et al. (20) defines two paradigms for designing RNA structures, namely, the positive and negative design paradigms. While positive design seeks a sequence with high affinity for the target structure, negative design aims for a sequence exhibiting high specificity to the target structure. To be successful, RNA design algorithms should preferably satisfy the objectives of both paradigms.

Various algorithms have been implemented to facilitate the search for RNA sequences that fulfill the design criteria [review: (19)], a list of implementations is available in Supplementary Table 1. Many methods allow the user to specify additional constraints or restraints, such as specific GC content, the presence of specific sequence or structure motifs, and the absence of forbidden patterns. Some of the recently developed methods are geared towards more specific problems, such as designing RNA sequences that fold into multiple target structures [e.g. RNAdesign (21)] or designing RNA sequences in the context of a specific 3D structure (22).

An experiment known as Eterna has been established to challenge computer algorithms and human participants with various design puzzles (23). It has provided a comparative benchmark and provided insight into structural features that govern the “designability” of individual RNA structures as well as of structural switches.

Considering the existing challenges in this field, we have developed a new tool for structure-based RNA design. DesiRNA explores the RNA sequence structure landscape using the Replica Exchange Monte Carlo (REMC) approach with the goal of generating a sequence that folds into the user-defined shape and also satisfies additional constraints. It allows the identification of sequences that fold into more than one desired structure, and it controls the ability of RNA to oligomerize. Our method combines the principles of positive and negative design and ensures that the generated sequences not only have a high affinity for the target structure but also high specificity that prevents folding into non-target structures.

## MATERIALS AND METHODS

### Design algorithm

Figure 1 depicts the basis of the DesiRNA algorithm.

**Figure 1:**
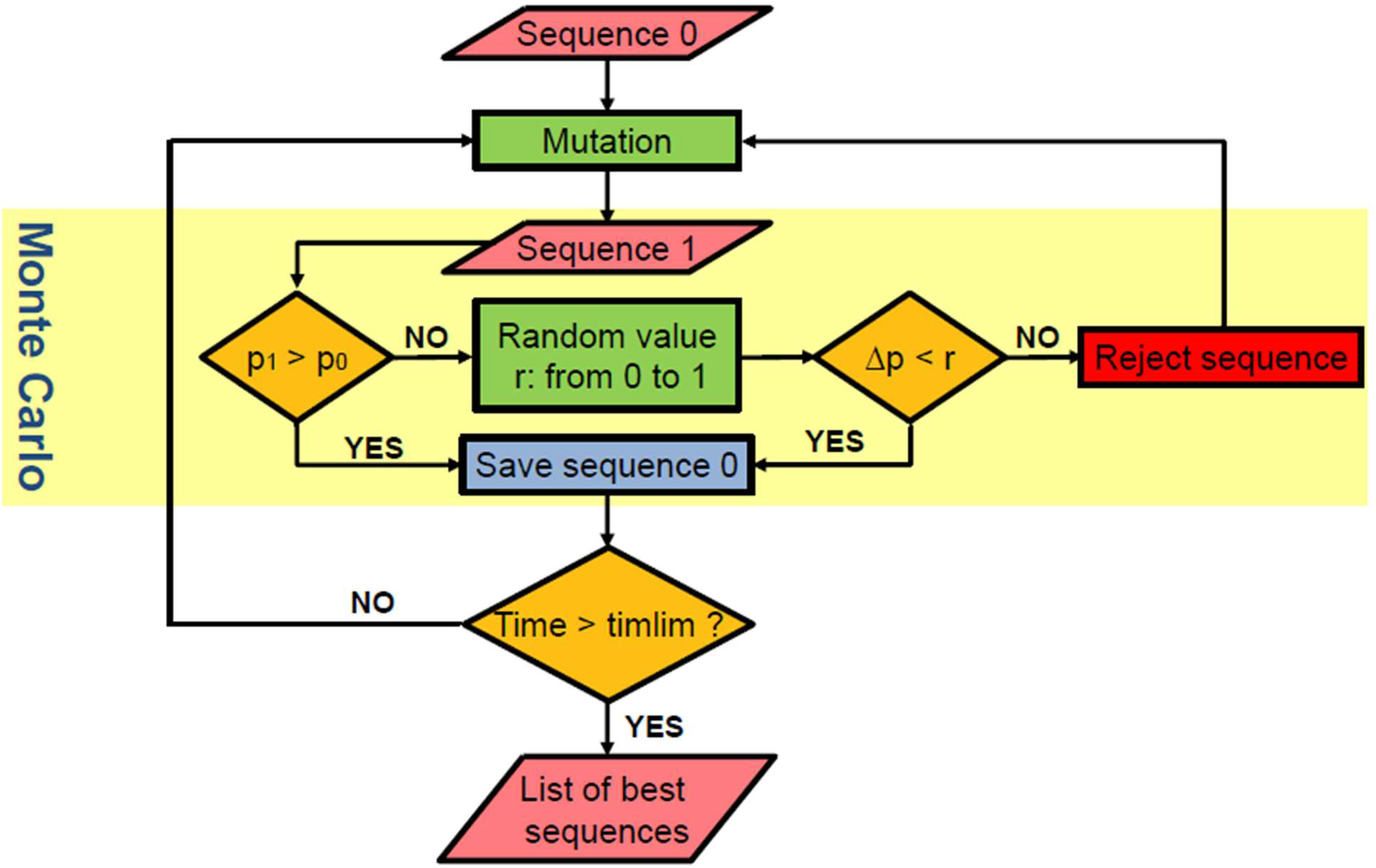
A workflow of the DesiRNA algorithm.

1. DesiRNA starts the RNA sequence design process with the generation of an initial sequence in accordance with the input structural and sequence constraints. This sequence generation follows the Eterna rules for RNA folding, providing an effective starting point for the algorithm.
2. The sequence is subjected to the computational prediction of the structure corresponding to the minimum free energy (MFE) conformation, and the corresponding energy and the base-pairing patterns are recorded. The energy of the fit of the sequence to the target structure is calculated, and the difference between the two energy values is recorded. The discrepancy between the MFE structure and the target structure is also calculated. A score is calculated that combines: i) the similarity of the MFE secondary structure to the target secondary structure, ii) the evaluation of the difference between the MFE energy and the energy of the sequence forced to fold into the target structure, and iii) the satisfaction of the user-defined restraints.
3. Multiple replicas of the initial sequence are created, each assigned with a different Monte Carlo (MC) temperature value to guide the Replica Exchange (RE) operations.
4a. Each replica sequence is subjected to an attempted mutation. The mutation point is selected randomly, and the mutation is either a single nucleotide change for an unpaired nucleotide or a double mutation for a pair of nucleotide residues. For each attempted mutation, the score is calculated as in point 2.
4b. If the score of the sequence with an attempted mutation is better than the score of the original sequence in a given replica, the mutation is accepted and replaces the pre-mutation sequence (along with its score) in the replica. If the score after the mutation is is equal to or smaller than the score before the mutation, the Metropolis criterion is applied.
4c. The Metropolis criterion involves the calculation of a probability distribution (p) based on the difference of the score differences for the mutated and original sequences, considering the temperature of the replica. A random number (n) between 0 and 1 is generated. If p exceeds n, the mutated sequence is accepted and replaces the previous sequence. Otherwise, it is rejected, and the original sequence is retained. Steps 4a-c are carried out for all replicas in a single step of the sequence design process.
5. After a defined number of steps, an attempt is made to exchange sequences between the replicas in the neighboring temperature shelves. The acceptance or rejection of which is based on the Metropolis criterion, depending on the score difference and the temperatures of the replicas.
6. The simulation continues for the specified number of steps, generating RNA sequences that attempt to simultaneously satisfy the following three goals: i) the MFE structure for the designed sequence is identical (or at least very similar) to the target structure, ii) the energy of the folded sequence is maximally lower than that of the various alternative structures that differ from the target structure, and iii) the user-defined restraints are satisfied as much as possible.

### Constraints and restraints in DesiRNA

Constraints are conditions that the solution must strictly adhere to. They set hard limits on the possible outcomes and cannot be violated. In RNA sequence design with DesiRNA, the constraints include: i) specific sequence and ii) specific base pairs. The user can constrain the sequences to be generated in the simulation to have a specific residue such as A or a subset of residues such as G or C at a specific position. If a base pair is specified for any two nucleotides, only sequences that have residue duplications compatible with that base pair can be generated, including the canonical pairs A-U, G-C, or the wobble pair G-U. If the user specifies constraints that are inherently inconsistent (e.g., that a base pair must be formed from residues at specific positions and these residues must be simultaneously retained as A and G), DesiRNA will inform the user that a solution that satisfies the constraints cannot be obtained.

Restraints are the desired parameters of the solution, and the conformance of the solution to the restriction is an element of the evaluation function. Restraints steer the design in a certain direction, but also allow for deviations. DesiRNA can optimize the conformance of the solution to the restraints, even if they cannot be fully satisfied. For example, a constraint could be included to specify a low GC content that is inconsistent with the constraints, such as a large number of G residues in the sequence contained in base pairs. Violation of restraints reduces the score of a solution, but does not prevent DesiRNA from considering that solution.

Importantly, DesiRNA considers the desired secondary structure as a restraint, while the possibility of forming base pairs by single residue doublets in the structure is considered a constraint. Therefore, the output of DesiRNA may contain sequences that fold into a structure that is very close to, but not completely identical to, the target structure, while optimizing compliance with all restraints and satisfying all constraints.

As input, DesiRNA requires a file with sequence and base pairing constraints that define the target secondary structure. Optionally, DesiRNA can accept additional restraints. The sequence constraints are specified in terms of IUPAC codes that denote different types of RNA nucleotides. For example, “N” represents any nucleotide, “Y” represents pyrimidines (C or U), “S” refers to G or C, etc. Secondary structure constraints are described in the commonly used dot-bracket notation, with paired nucleotides indicated by matching brackets “()” and unpaired nucleotides indicated by dots “.”.

The DesiRNA output is a CSV-formatted table listing the highest-scoring sequences (10 by default). Each row indicates the secondary structure, the corresponding sequence, a relative score (from 0 to 1 - the higher, the better), and the percent residue (AUCG) of the sequence.

### Secondary structure prediction and energy evaluation

In DesiRNA, RNA secondary structure prediction and energy evaluation can be done essentially by any method that can be plugged into the Python framework. The default option is to use the ViennaRNA package (34). In particular, the ‘pf_fold‘ function from ViennaRNA is used to calculate the minimum free energy (MFE) of an RNA sequence and the ‘energy_of_structure‘ function is used to calculate the energy of an RNA sequence folded into the target secondary structure. To assess the oligomerization potential of the sequences, the ‘mfe‘ method from the ‘fold_compound‘ class is used to calculate the MFE of a given sequence in a monomeric vs a dimeric state.

When designing RNA structures with pseudoknots, DesiRNA always generates sequences compatible with the formation of all desired base pairs. However, evaluating the structures that can be generated by this sequence and evaluating the energy of a fit to the structure with pseudoknots requires the use of a method that accounts for pseudoknots, such as IPknot (11). This approach can make DesiRNA simulation more computationally expensive and thus more time consuming. We developed an alternative solution that can be considered as a workaround that allows the consideration of pseudoknots as a modification of the original predictions without pseudoknots. In the default option used by ViennaRNA, DesiRNA first folds the structure or computes the energy of the fit to the structure without the pseudoknots. In the next step, additional non-nested helices contributing to the pseudoknots are introduced, and the energy of the affected parts of the sequence is recalculated using Turner’s thermodynamic parameters.

### Design of sequences compatible with multiple alternative structures

DesiRNA’s multi-structure design strategy uses ViennaRNA’s ‘‘subopt’’ functionality. The user specifies structural constraints for, say, two different structures that a single sequence could fold into. ViennaRNA’s ‘‘subopt’’ functionality is then used to identify a set of suboptimal secondary structures within a given energy range of the minimum free energy structure. DesiRNA then attempts to identify a sequence that maximizes the match of the predicted MFE conformation to one of the target structures, as well as to obtain an alternative suboptimal structure with a low energy range with maximum similarity to the other target structure. With this approach, the design is directed toward sequences with inherent structural flexibility, such as those that can adopt two predetermined conformations. This approach can be used to design riboswitches or other functional RNA molecules that function by transitioning between different structural states.

### Availability

DesiRNA is freely available on GitHub (https://github.com/fryzjergda/DesiRNA.git). The software is implemented in Python 3 and includes a comprehensive installation manual providing detailed instructions for setting up and configuring all required libraries and dependencies.

## RESULTS

We developed DesiRNA, a program for structure-based RNA sequence design that generates RNA sequences that, under defined constraints, are predisposed to fold into specific secondary structures and fulfill user-defined restraints. Its attributes include the ability to handle pseudoknotted structures and the capacity to assess the oligomerization propensity of the designed sequences, which can be used to design RNA-RNA complexes or to promote or prevent the formation of oligomers. The tool also supports the design of sequences capable of folding into multiple target structures. To enable the design of RNA sequences that resemble natural molecules (e.g., without homopolymeric stretches or repetitive motifs), DesiRNA enables the control of GC base pair percentage. Its approach to design is bifunctional: In the positive design paradigm, it employs partition function calculations to optimize the likelihood of the designed sequence adopting the target structure. In the negative design paradigm, the algorithm seeks to maximize the energy difference between the minimum free energy (MFE) structure of the designed sequence and the alternative conformations with suboptimal energies.

### DesiRNA benchmark

The Eterna puzzles (23) is a collection of 100 RNA secondary structure patterns, known to provide a difficult test for RNA design algorithms. For the purpose of tests presented in this version of the article, a puzzle was deemed solved if the algorithm was able to produce a solution within the allocated time constraint. The current state-of-the-art in the field is represented by MoiRNAiFold (24), which has solved 91 of the Eterna puzzles within the 24-hour limit.

The current version of DesiRNA managed to solve 82 puzzles within the 24-hour limit. To the best of our knowledge, DesiRNA is the only fully automated method that was able to solve puzzles number 78, 89, and 97. These specific puzzles have proven to be formidable challenges for other algorithms, highlighting the unique capabilities of DesiRNA in the context of RNA design.

Figure 2 shows a comparative overview of the ability of a number of RNA design algorithms to solve Eterna puzzles.

**Figure 2:**
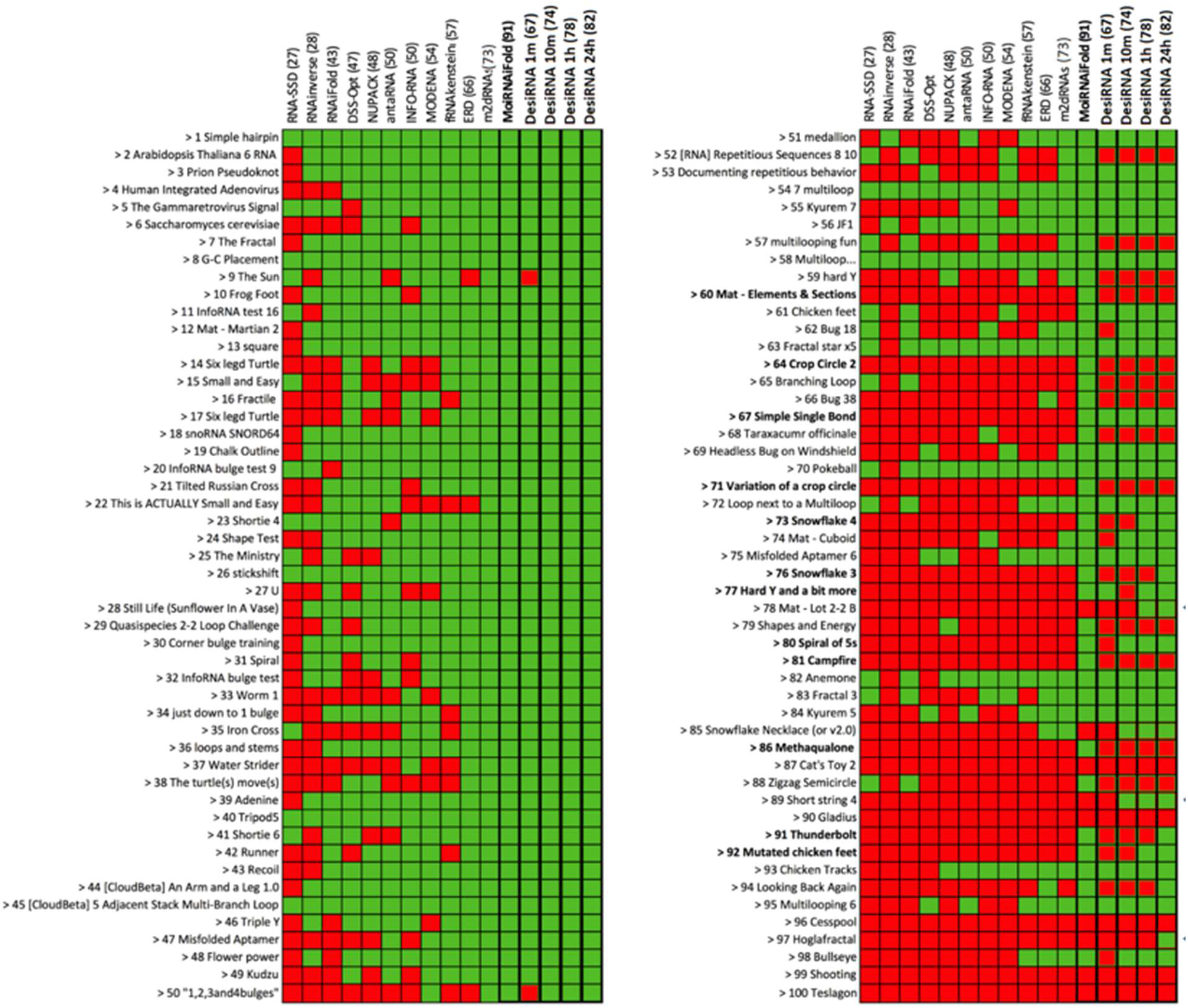
DesiRNA benchmarking results and comparison with published RNA sequence design tools in the field. The green color indicates the successful solution of the RNA design problem, and the red color indicates the lack of the ideal solution. Results for DesiRNA indicate the performance after 1 minute, 10 minutes, 10 hours, and 24 hours, respectively.

## DISCUSSION

To address the challenges of RNA sequence design, we have developed DesiRNA, a method to design RNA sequences that fold into specific target structures while accommodating a set of user-defined constraints. DesiRNA was specifically designed to support the design of sequences that resemble natural RNA sequences, fold into functional 3D structures, contain features typical of functional RNAs such as pseudoknots, account for oligomerization, and most importantly, retain the sequence composition typical of natural RNAs. An important feature of DesiRNA is the ability to design RNA sequences that can fold into multiple structures, allowing the design of RNAs that are able to switch between functional states in response to environmental cues.

We have subjected the current version of DesiRNA to evaluation using the Eterna benchmark. The results provide a comparative analysis of DesiRNA’s performance against other RNA design algorithms, highlighting both its strengths and weaknesses. The current version of DesiRNA solved 67 puzzles in a one-minute run, a number that gradually increased to 74 in 10 minutes, 78 in an hour, and 82 in a 24-hour period. For the current version of our method, this performance represents the second best result in the field, surpassed only by MoiRNAiFold.

Most importantly, DesiRNA was able to solve puzzles 78, 89, and 97. These puzzles pose particular challenges in the field of RNA design, as they are usually problematic due to their complicated structures.

## ACKNOWLEDGEMENTS

This work was funded by the Polish National Science Center (NCN), grant number: 2017/25/B/NZ2/01294 to J.M.B.

